# Performance of the 10X Genomics Flex Single-Cell Sequencing Assay and its Application to Overcome Challenges in Clinical Trial Samples

**DOI:** 10.64898/2026.03.06.710062

**Authors:** Martina Antoniolli, Llucia Albertí Servera, Kerstin Paetzold, Stephan Schmeing, Carmen Yong, Sina Nassiri, Tamara Hüsser, Michael A. Cannarile, Marina Bacac, Emilio Yángüez, Steffen Dettling

## Abstract

**Background:** Single-cell RNA sequencing (scRNA-seq) has become essential for understanding disease biology, yet its application in clinical trials is often limited by logistical challenges associated with the handling of biospecimens. Newly developed protocols aim to address these limitations by enabling profiling of fixed tissue. These new solutions need to be benchmarked against well-established protocols to assess their performance and suitability for clinical research.

**Results:** We systematically compared different scRNA-seq protocols in a set of samples commonly analysed in clinical trials. The 10X Genomics GEM-X Flex Gene Expression assay (GEM-X Flex), in combination with the “chop-fix” preprocessing protocol, demonstrated superior performance to the standard GEM-X Universal Gene Expression solution when applied to both primary tumor tissue fragments and FFPE blocks. Moreover, the quality of the data obtained from GEM-X Flex applied to FFPE blocks outperformed that of single-nuclei RNA sequencing (snRNA-seq) from frozen biopsies, more robustly capturing the biological signals associated with the mechanism of action of a drug evaluated in an internal clinical trial.

**Conclusions:** GEM-X Flex generates reliable, comprehensive transcriptomic data from both fixed tissue and clinical biopsies. By overcoming some of the limitations of fresh and frozen tissue analysis, this protocol offers a robust solution for the broad implementation of scRNA-seq in clinical trials.

## Introduction

The development of single-cell RNA sequencing (scRNA-seq) has revolutionized molecular biology, enabling detailed insights into cellular heterogeneity and gene expression dynamics across diverse biological contexts. In the past fifteen years, the development of sensitive protocols^1–3^ combined with efficient solutions for cell compartmentalization and barcoding^4–7^ have exponentially increased the throughput of single-cell analysis^8^. Moreover, the commercialization of reliable single-cell solutions from different vendors has significantly facilitated the access to single-cell technologies and their broad applications to different fields^9–12^. Under the umbrella of the Human Cell Atlas (HCA) consortium, these efforts have contributed to a comprehensive map of human cells, significantly advancing our understanding of human biology and disease^13–15^.

Despite these advancements, the general application of scRNA-seq in clinical trials remains limited due to several challenges^16^. As a result, most clinical applications of genomics rely on bulk approaches, which do not capture the variation underlying cellular heterogeneity. High costs and technical hurdles, particularly in the handling of fresh clinical specimens such as core needle biopsies, continue to hamper scRNA-seq implementation^17^. One of the primary issues when working with fresh material is the time sensitivity associated with sample processing^18^. Successful scRNA-seq requires high-quality samples with sufficient numbers of viable cells. However, in multi-center clinical trials, where the medical teams are primarily focused on patient care, it is often difficult to preserve tissue specimens in a standardized way^19^. The separation of sample collection and processing teams, along with limited access to necessary preservation materials, further complicates the process. The unpredictable nature of operating theaters or outpatient clinics makes it difficult to ensure standardized preservation protocols, increasing the risk of RNA degradation and compromising sample quality, ultimately affecting the quality of subsequent biological insights^20^.

To overcome some of these challenges, clinical specimens can be frozen on-site and shipped to centralized labs for scRNA-seq analysis. However, this procedure may still pose logistical challenges, especially at a clinical site that lacks the infrastructure to quickly freeze samples. Single-nucleus RNA sequencing (snRNA-seq) has emerged as a reliable alternative for profiling frozen and difficult-to-dissociate samples, such as core needle biopsies^10,20^. This method decouples tissue acquisition from immediate sample processing, allowing the analysis of challenging clinical specimens. However, snRNA-seq has its own limitations, including lower mRNA content in nuclei compared to whole cells, skewed transcript representation, and the impossibility of enriching specific cell types of interest based on cell surface markers^21,22^. Furthermore, ambient RNA contamination is a significant issue during nuclei preparation, which requires additional filtering steps that can compromise biological insights derived from the data^10^. This issue is especially problematic in clinical trials, where consistency and reproducibility are critical.

In the past years, fixed tissue samples, particularly formalin-fixed paraffin-embedded (FFPE) blocks, have gained increasing attention as an alternative to fresh or frozen tissue in clinical trials^23^. To enable the analysis of such fixed samples, highly sensitive protocols for transcriptomic and spatial analysis have been developed^24–33^. Following this idea, different groups have recently focused on the development of single-cell protocols in which fixed tissue can be used as starting material^34–41^. These efforts have consolidated in the introduction of a commercial solution that allows performing scRNA-seq at a large-scale from fixed tissue^42,43^. These advancements hold promise of overcoming some of the above-described limitations associated with the implementation of single-cell analysis in clinical trials.

In this study, we aimed to evaluate the performance of the 10X Genomics GEM-X Flex Gene Expression (GEM-X Flex) assay on clinical samples, benchmarking it against existing solutions from 10X Genomics. We compared different preservation and pre-processing strategies to determine which workflow is best suited for clinical settings. Finally, we evaluated the performance of the GEM-X Flex protocol on FFPE blocks from an internal Phase I dose escalation study evaluating RO7119929 (clinical trial NCT04338685)^44^. We compared the data quality and the biological insights derived from the fixed FFPE blocks with those obtained from frozen core needle biopsies from the same patients as part of the clinical trial. Our findings will inform the selection of optimal protocols for using scRNA-seq in clinical studies, helping to choose the most appropriate sample preservation method and processing strategy for precious clinical specimens.

## Results

### Initial benchmarking of scRNA-seq workflows for the analysis of tumor samples

To identify the optimal workflow for analyzing clinical tumor samples, we systematically compared two commercial kits released by 10X Genomics: the GEM-X Universal 5’ Gene Expression and the GEM-X Flex Gene Expression, the latter tested in combination with different pre-processing strategies (**Figure 1A**). As a surrogate for core-needle biopsies and to limit the confounding effects of inter-patient variability, we evaluated each protocol’s performance using three randomized tumor fragments generated from a commercially available primary lung cancer sample. Using tumor-derived fresh nuclei suspensions as input material, half of the sample was processed with the well-established 10X GEM-X Universal 5’ Gene Expression protocol (sample **A**), while the other half was fixed and processed according to the new GEM-X Flex protocol (sample **B**). Moreover, to recreate the storage conditions most frequently used in our clinical studies, we snap-froze additional tumor fragments and subsequently isolated and fixed the nuclei suspension using the GEM-X Flex chemistry (sample **C**). Finally, we directly fixed the remaining tumor fragments and processed them according to the published 10X Genomics Chop/Fix protocol. Here, the tumor tissue is first fixed and stored and then dissociated into single-cell suspension prior to the use of 10X Genomics GEM-X Flex protocol (sample **D**).

**Figure 1.**
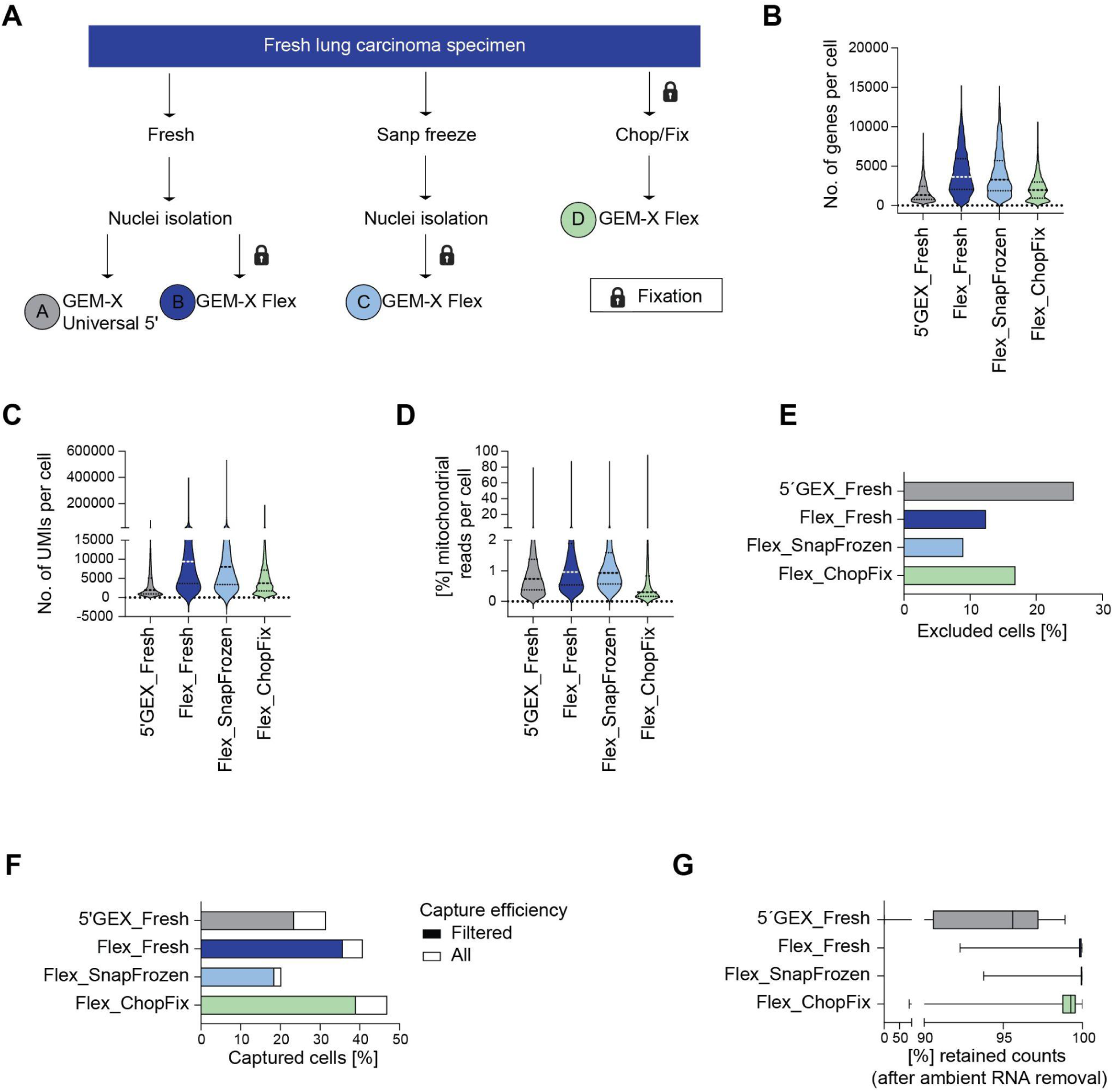
Experimental workflow for the tumor samples and initial Quality Control (QC) metrics. **(A)** Diagram for experimental conditions. Fresh tumor specimens from a primary lung carcinoma sample were manually dissected, randomized, and processed according to the protocols represented in the figure. **(B-D)** Distribution of the number of genes **(B)**, the number of Unique Molecular Identifiers (UMIs, **C**), or the percentage of reads mapping to mitochondrial genes **(D)** per nuclei/cell detected in the samples processed with the different protocols (non-filtered CellRanger cells, only genes included in the GEM-X Flex probeset). **(E)** Percentage of nuclei/cells excluded per sample due to low quality, based on the thresholding applied to the above-mentioned quality metrics together with the CellBender filtering as explained in Material & Methods. **(F)** Capture efficiency for the different samples defined as the percentage of loaded nuclei/cells that were initially sequenced according to CellRanger (outer boundary of the bar, All) or that passed QC (filled with color, Filtered). **(G)** Distribution of the percentage of retained counts per nuclei/cell after ambient RNA removal using CellBender in the different samples (only nuclei/cells that passed QC are represented).

We initially generated basic library QC metrics and compared the number of genes (**Figure 1B**) and unique molecular identifiers (UMIs) (**Figure 1C**) detected per sequenced cells. Both the number of genes and UMIs per cell in the sample processed with the 5’ chemistry were lower than in the ones processed with the GEM-X Flex chemistry. These observations are true even if all genes, rather than just the ones included in the GEM-X Flex probeset, were considered in the analysis of the sample processed with the 5’ chemistry (**Supplementary Figure 1 A-D**), suggesting a superior sensitivity of the GEM-X Flex. When focusing on the fixed samples processed with the GEM-X Flex, the Chop/Fix protocol captured fewer genes and counts than its counterparts, a trend that persisted even after read downsampling and is not caused by insufficient sequencing depth as shown by the CellRanger analysis plots (**Supplementary Figure 1E**). We next focused on the analysis of percent of reads mapping to mitochondrial genes and, as expected for nuclei samples, all conditions exhibited a low percentage of mitochondrial reads (**Figure 1D**). Importantly, mitochondrial percentages across different chemistries are not directly comparable. However, among the samples processed with the Flex technology, Chop/Fix libraries contained the lowest mitochondrial read percentage, with a median of 0.3% compared to approximately 0.9% in the other workflows (**Supplementary Figure 1D**).

After applying a cut-off for low quality cells based on the QC metrics described above, more than 70% of the cells were retained in all cases (**Figure 1E**). In the sample processed with the 5’ chemistry, the highest number of cells were filtered out of the analysis (25.68%), followed by the sample processed with the Chop/Fix protocol (16.87%) and the two nuclei samples processed with the Flex chemistry (12.39% and 8.96% respectively). Regarding the total capture efficiency, defined as the ratio between the retained cells after filtering and the number of cells loaded into the Chromium controller, the Chop/Fix protocol was superior to the rest of the workflows (**Figure 1F**). However, we cannot completely rule out that these differences are mostly driven by inaccuracies in cell counting before loading the chip. Furthermore, when applying ambient RNA correction using CellBender, the samples processed with the Flex kit retained more counts than the 5’ sample, suggesting a higher presence of ambient RNA in the last one (**Figure 1G and Supplementary Figure 1F**). Altogether, the general data quality and the sensitivity in the samples processed with GEM-X Flex chemistry is slightly superior to those processed with the 5’ chemistry.

### Cell type abundance differs across the tested single-cell sequencing protocols

The ability of different methods to recover and resolve cell types varies due to factors like capture and amplification efficiency, but also cell characteristics such as size, robustness, and mRNA content. Efficient recovery across diverse cell types is crucial for accurate analysis, especially in low-quality tissues or samples with a limited number of cells. To evaluate the performance of the different workflows in recovering diverse cell types, the cellular composition across samples was analyzed. Overall, we profiled 4.683 cells with the GEM-X Universal 5’ Gene Expression and 17.068 cells with the three GEM-X Flex protocols. Initially, UMAP analysis with ambient RNA correction, probeset restricted genes, but no integration methods (**Supplementary Figure 2 A,B**), revealed that the samples clustered based on the technology used. The samples processed with the 5’ chemistry and those processed using the GEM-X Flex chemistry occupied separate spaces in the UMAP plot. This indicates that without integration, the differences between the chemistries overshadowed the biological similarities of the samples. To address these batch effects, we applied Batch-Balanced K-Nearest Neighbors (BBKNN) integration, which significantly improved the clustering of the cells by their cell type and resulted in good sample representation across the various clusters (**Figure 2A**).

**Figure 2.**
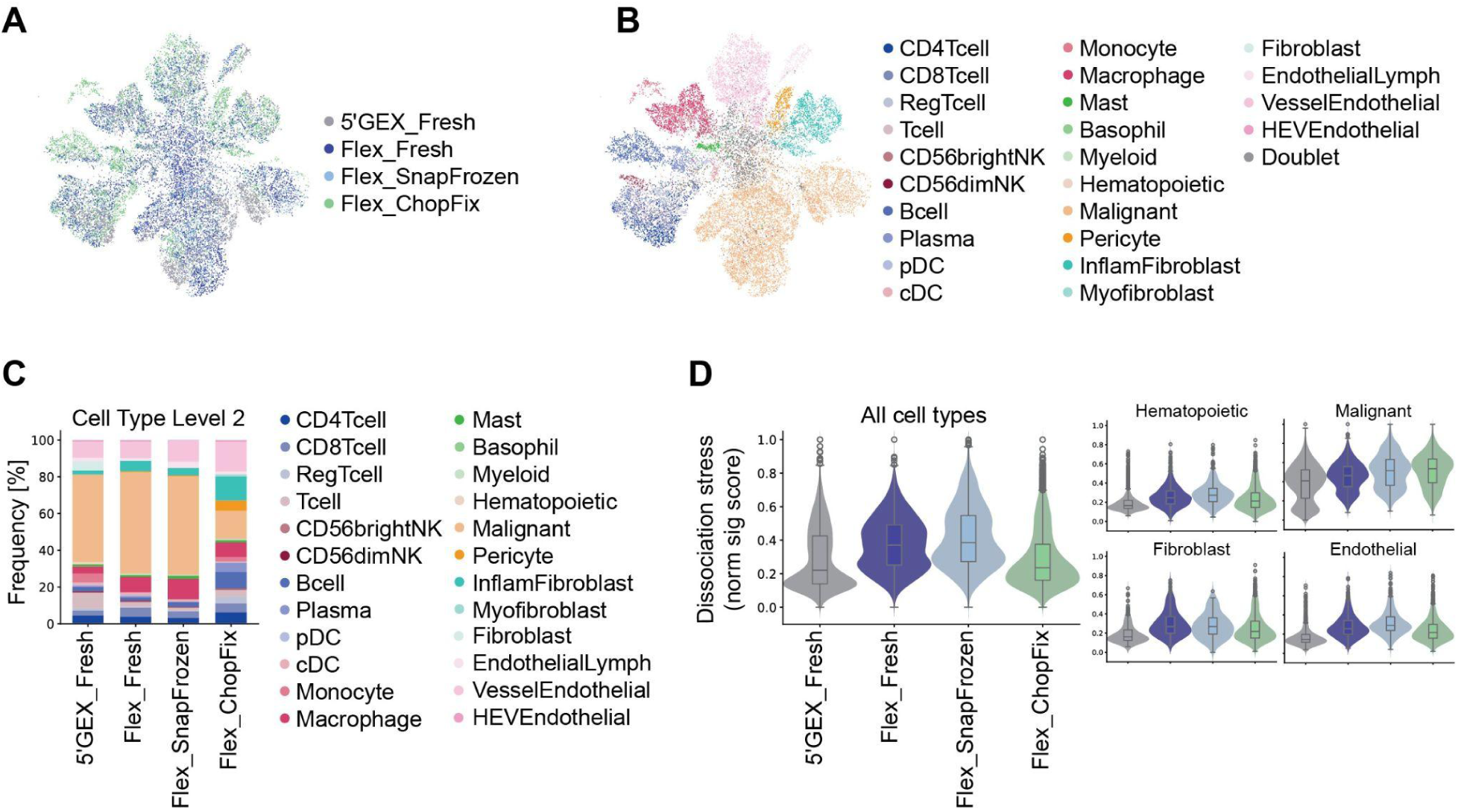
Sample clustering, cell type annotation, and quantification from the tumor samples. **(A)** Uniform manifold approximation and projection (UMAP) plot of data after Batch Balanced K-Nearest Neighbors (BBKNN) integration, colored by experimental condition. **(B)** UMAP plot of data after BBKNN integration, colored by cell type (granular annotation level 2). **(C)** Cell type distribution as percentage of the total number of nuclei/cells identified in the different samples (granular annotation level 2). **(D)** Dissociation stress is represented as a normalized gene signature score (min=0, max=1) calculated per sample and shown either for all cell types or for selected relevant cell types (Hematopoietic, Malignant, Fibroblast, Endothelial cells).

We successfully identified the main cell types anticipated in the sample (**Figure 2B, Supplementary Figure 2C**), such as various immune cell subsets, fibroblasts, and endothelial cells, across the different technologies. When we quantified the distribution of cell types across the different samples, we noticed that samples initially processed through nuclei generation exhibited a more similar cell type distribution. The sample processed with the Chop/Fix protocol displayed a higher proportion of fibroblasts and hematopoietic cells, predominantly B- and T-cells, and a reduced proportion of malignant cells compared to the other samples (**Supplementary Figure 2 D,E**). Additionally, we observed an increased percentage of doublets in the nuclei samples processed with Flex chemistry (**Supplementary Figure 2F**), which may have arisen from nuclei crosslinking during PFA fixation before loading onto the 10X Genomics Chromium controller. By examining the data with greater granularity, we encountered limitations in resolving certain cell types with the different protocols (**Figure 2C**). In the sample processed with the 5’ chemistry, some fibroblasts, T-cells, and general hematopoietic cells could not be further dissected. In samples processed with Flex chemistry, most of the cells could be resolved except for some T and myeloid cells.

To further explore whether the different protocols may induce dissociation-related transcriptomic changes, we evaluated a gene signature associated with cellular stress responses as previously described^45,46^. The normalized stress score has a slightly higher proportion of cells with lower scores in the Chop/Fix and 5’ chemistry samples (**Figure 2D**). However, we observed greater differences in the stress score distribution between different cell types, with malignant cells having more cells in the higher score part than hematopoietic, fibroblasts, and endothelial cells. These differences can be explained by the fact that malignant cells are especially susceptible to *ex-vivo* manipulation. Altogether, the cell type abundances slightly differ across the tested single-cell sequencing protocols, with less abundant and more fragile cell types being better captured in the samples processed with the Chop/Fix protocol.

### The GEM-X Flex protocol generates high-quality single-cell data from clinical FFPE blocks

The GEM-X Flex technology additionally allows performing scRNA-seq from FFPE blocks. To investigate the performance of GEM-X Flex technology applied to clinical trial samples, we retrospectively compared data generated from two different scRNA-seq workflows. In particular, we analyzed nuclei suspensions obtained from slow-frozen needle core biopsies, processed with the standard 10X Genomics 3’ chemistry as part of RO7119929 clinical trial (NCT04338685)^44^, and benchmarked them against newly generated scRNA-seq data obtained from FFPE blocks from three matching patients processed with the GEM-X Flex protocol (**Figure 3A**). For both sets, we analyzed baseline (untreated) and RO7119929-treated (on-treatment) samples, allowing us to compare how the different technologies are able to capture RO7119929-induced transcriptional changes and specific gene signatures associated with its mode of action.

**Figure 3.**
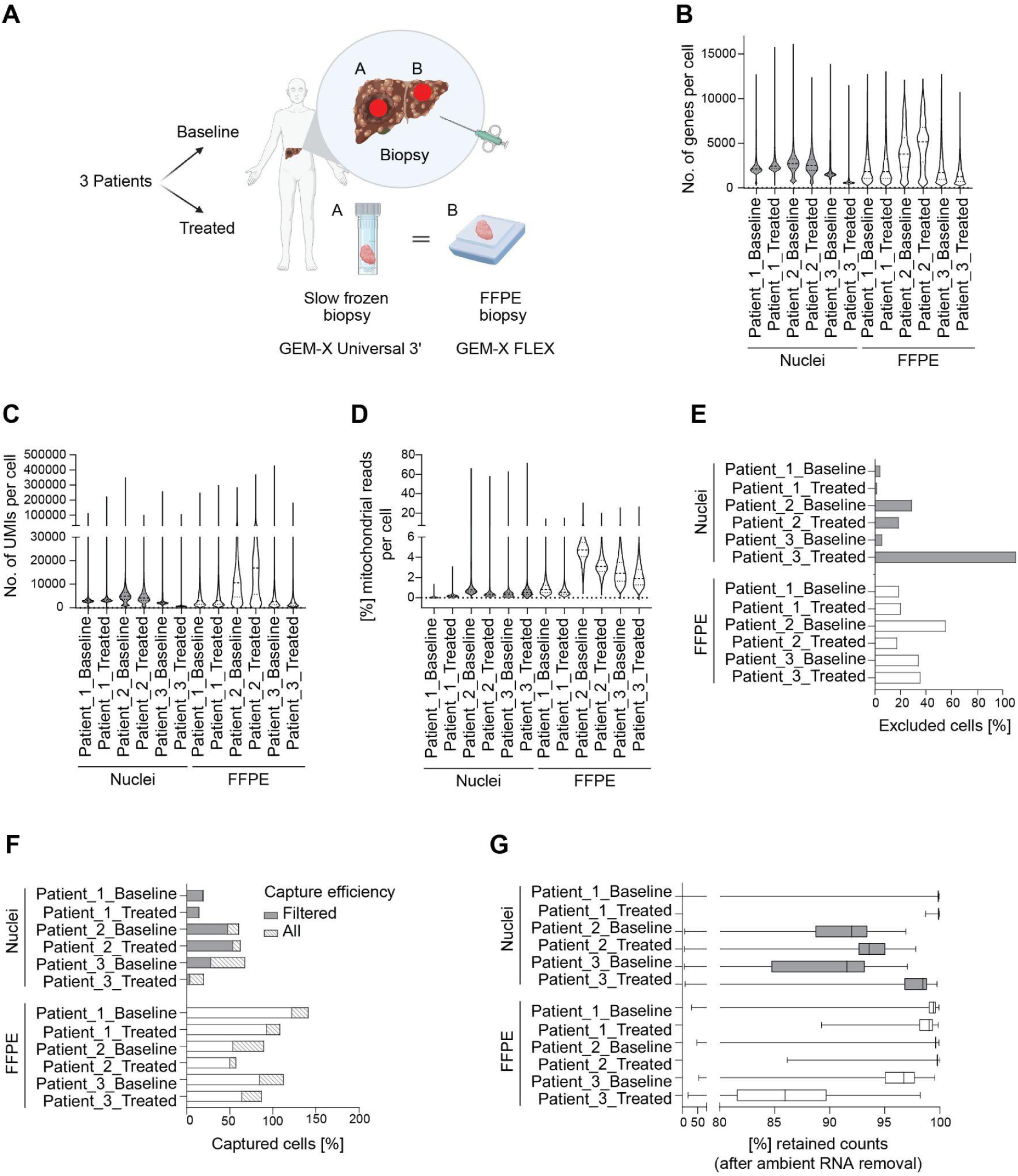
Experimental layout for clinical samples and initial quality control metrics. **(A)** As part of the NCT04338685 clinical trial, slow-frozen tumor biopsies collected from three patients (both at baseline and on-treatment) were analyzed by snRNA-seq using the Chromium Single Cell 3ʹ v3.1 protocol. Matching Formalin-Fixed Paraffin-Embedded (FFPE) biopsies were retrospectively processed with the 10X Genomics GEM-X Flex Gene Expression protocol. **(B-D)** Distribution of the number of genes **(B)**, the number of Unique Molecular Identifiers (UMIs, **C**), or the percentage of reads mapping to mitochondrial genes **(D)** per nuclei/cell detected across all three patient samples (Baseline and Treated) and processed with the different protocols. Included non-filtered CellRanger cells and all detected genes. **(E)** Percentage of nuclei/cells per sample excluded due to low quality, based on thresholding applied to the above-mentioned QC metrics together with the CellBender filtering as explained in Material & Methods. **(F)** Capture efficiency for the different samples defined as the percentage of loaded nuclei/cells that were initially sequenced according to CellRanger (outer boundary of the bar, All) or that passed QC (filled with color, Filtered). **(G)** Distribution of the number of counts per nuclei/cells retained after ambient RNA removal using CellBender in the different samples (only nuclei/cells that passed QC are represented).

First, we aimed to test the suitability of the GEM-X Flex protocol for clinical trial samples, by comparing the data quality focusing on general QC metrics. Overall, a slightly higher number of genes and UMI counts were detected in the FFPE samples compared to nuclei samples (**Figure 3 B,C**). All samples contained a low number of mitochondrial reads for their respective chemistries, with the nuclei samples exhibiting less than 1% and FFPE samples showing less than 5% of mitochondrial RNA (**Figure 3D)**.

After applying a cut-off for low-quality cells, more than 60% of the cells were retained in all the biopsies (**Figure 3E**), except the nuclei on-treatment sample from Patient 3. Here, 79.48% of cells were discarded, failing to meet our QC threshold and therefore excluded from the second part of the analysis. Regarding the total capture efficiency, the samples processed with the Flex protocol outperform the nuclei preparation (**Figure 3F**). However, this data may not be perfectly accurate, due to the challenges of counting cells derived from dissociated FFPE blocks. When performing ambient RNA correction (**Figure 3G**), in general, FFPE samples retained more than 95% of the counts, with the exception of the on-treatment sample from Patient 3. In contrast, half of the nuclei samples lost 5-10% counts when removing the signal attributed to ambient RNA contamination. In summary, we can conclude that the Flex protocol is effective in generating high-quality single-cell data from clinical FFPE blocks, improving the data quality usually obtained from frozen biopsies processed by snRNA-seq.

### The GEM-X Flex applied to FFPE blocks allows the identification of challenging cell types and states and improves the capture of relevant biological signals associated with drug mode of action

Next, we analyzed the cluster structure and cell type annotation on the different samples. When analyzing the data without using an integration method, the different samples cluster according to the processing protocol: nuclei samples derived from the frozen biopsies occupy a different space compared to the FFPE samples on the UMAP plot (**Supplementary Figure 3A**). Integration methods increase the sample overlap, and the different samples intermix (**Figure 4A**), with the cells clustering according to the cell type of origin (**Figure 4B**). The exception is the hepatocytes which, as expected, cluster per patient sample. We identified the different cell types expected in the liver biopsies (**Figure 4C**, and **Supplementary Figure 3B**), with hepatocytes being the major population, partially because they are almost the only cell type detected in both baseline and on-treatment biopsies from Patient 2. Focusing on the cell type distribution in the different samples, the FFPE blocks processed with the Flex protocol display a higher diversity, with more cell types represented especially at baseline.

**Figure 4.**
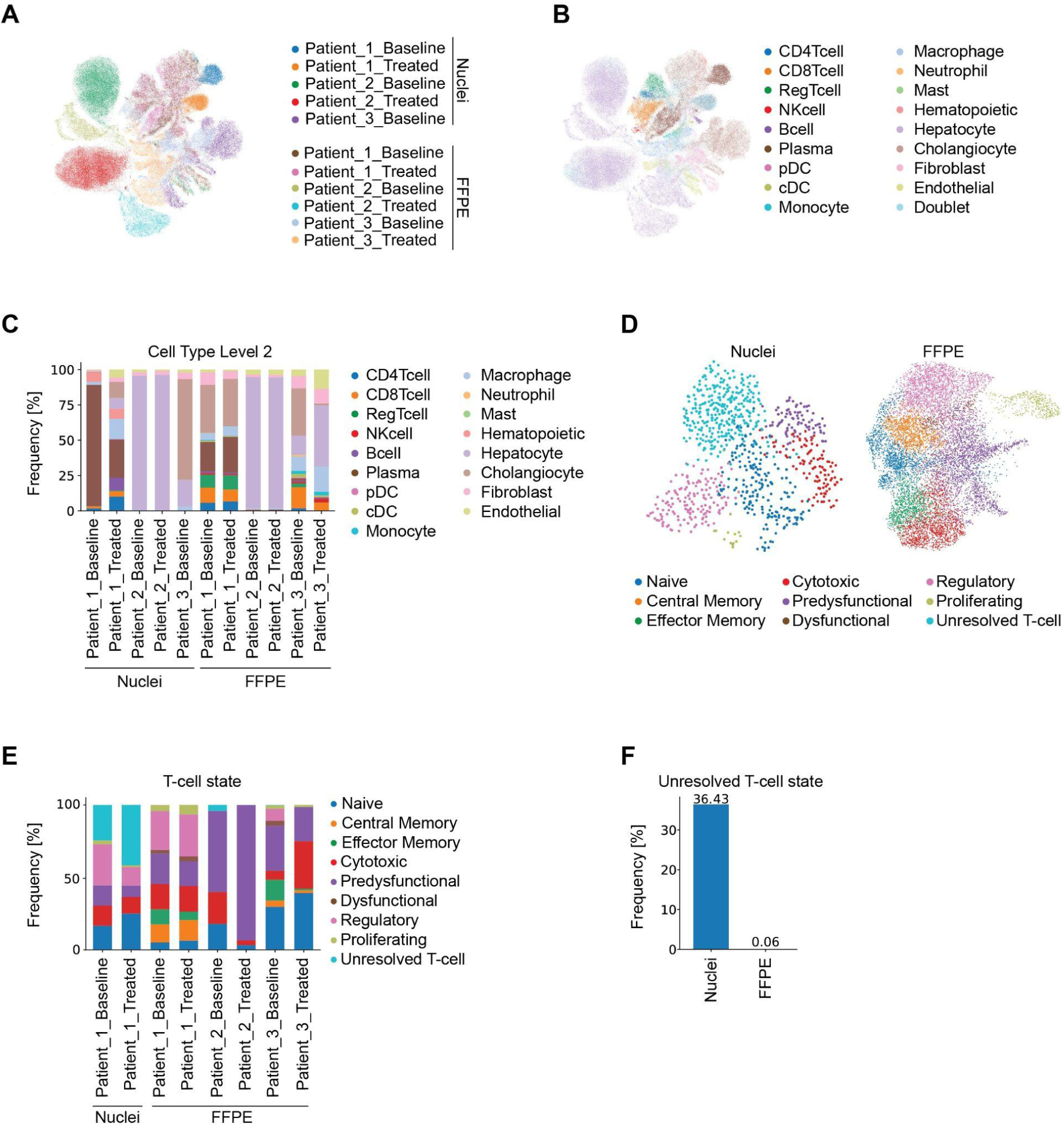
Sample clustering, cell type annotation, and quantification from clinical samples. **(A)** Uniform manifold approximation and projection (UMAP) plot of integrated data, colored by sample according to figure legend. **(B)** UMAP of integrated data, colored by cell type. **(C)** Cell type distribution as percentage of the total nuclei/cells identified in the different samples (granular annotation level 2). **(D)** UMAP plot of T-cells in samples derived from nuclei suspensions of slow-frozen biopsies (left) or FFPE blocks (right) colored according to T-cell states annotation. **(E)** T-cell states distribution as percentage of the total number of T-cells identified in the different samples. **(F)** Percentage of T-cells per processing protocol for which the T-cell state could not be resolved.

Next, we focused on the analysis of T-cells, which are key effectors of anti-tumor therapies such as check-point inhibitors or T-cell engagers, and compared how the processing protocols are able to resolve the different cell states. We identified 851 T-cells in the nuclei samples, coming exclusively from Patient 1, both at baseline and on-treatment. In contrast, we captured 11.698 T-cells in the FFPE samples from all three patients (**Figure 4D**). Using pre-defined gene signatures, T-cells were classified into different subpopulations in FFPE samples, while this was impossible when using the nuclei samples (**Figure 4 D,E**). Moreover, 36.43% of T-cells from the nuclei samples (**Figure 4D** upper left in UMAP plot) could not be resolved (**Figure 4F** and **Supplementary Figure 3C**) due to a mixed and very low signature score for the different states (**Supplementary Figure 3 D,E**). This data suggests that the GEM-X Flex protocol applied to FFPE blocks can help to resolve cell states that are otherwise more challenging to discriminate against when analyzing frozen tumor samples processed with the standard 10X Genomics 3’ chemistry.

Finally, based on the insights from the clinical trial, we compared the sensitivity of the different protocols to capture the RO7119929-induced increase of interferon stimulating genes, type1 interferon-related genes, and TLR7 downstream-related genes in liver tissue, associated with the expected mode of action of RO7119929^44^. We were able to detect and quantify the expression of *ISG15, MX1, XAF1, HERC5, STAT1* in all the samples analyzed (**Figure 5A**). In general, we observed bigger differences in expression between baseline and on-treatment samples in the FFPE blocks processed with the GEM-X Flex protocol, which resulted in higher fold-change values in these samples for all the genes in the panel with the exception of *MX1* (**Figure 5B**). Next, we analyzed macrophage polarization as an additional readout for TLR7+ responder cells upon exposure to the active form of RO7119929^44^. Using pre-defined gene signatures, we calculated the M1-like and M2-like macrophage scores in the different samples (**Figure 5 C,D**). In most of the samples, the treatment-induced change in average sample scores were higher in the FFPE samples processed with the GEM-X Flex protocol. This is especially true for the M1-like signatures, showing a re-polarization towards an M1-like immunostimulatory phenotype, which the standard 10X Genomics 3’ chemistry was not able to reliably capture. Altogether, the GEM-X Flex protocol not only allowed the identification of up-regulated biomarker genes upon treatment, but also revealed more pronounced treatment-induced changes of tumor-associated macrophages (TAMs) polarization when compared to the 3’ protocol.

**Figure 5.**
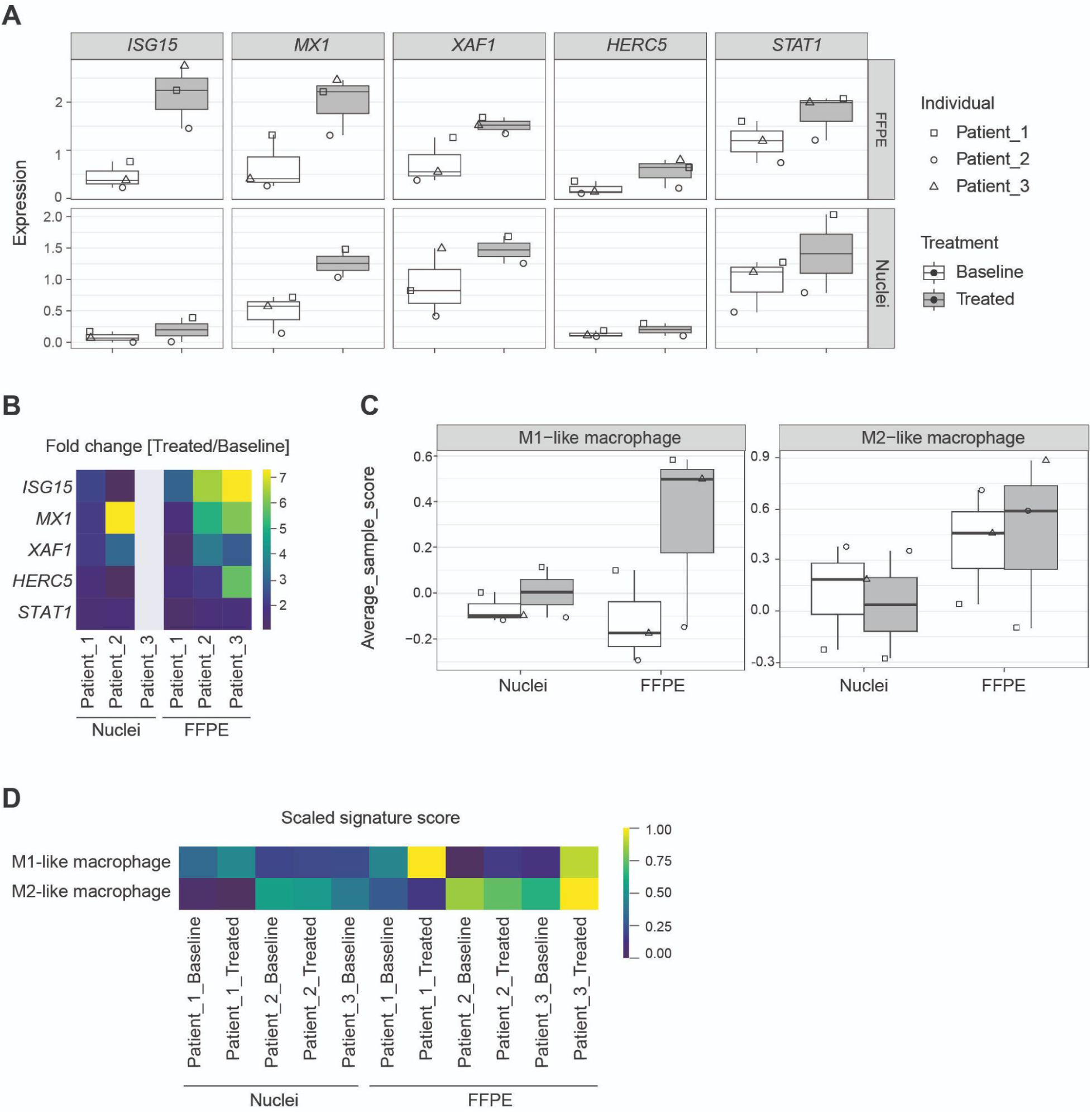
Identification of relevant gene signatures from the clinical study. **(A)** Gene expression analysis of *ISG15, MX1, XAF1, HERC5, STAT1* in Hepatocellular carcinoma displayed per patient (symbols), grouped per treatment condition (untreated: white; Treated: grey) and per sample of origin (top row FFPE, bottom row Nuclei). **(B)** Fold change (treated/baseline) revealed a strong induction of *ISG15* across all patients among PoMs which is more obvious in FFPE than in nuclei data FFPE samples. **(C)** Average macrophage state (M1 and M2) score per cell and chemistry. Data are displayed per patient (symbols), grouped per treatment condition (untreated: white; Treated: grey) and per sample of origin (top FFPE, bottom Nuclei). **(D)** Macrophage state (M1 and M2) score per sample and scaled per state.

## Discussion

Single-cell RNA sequencing has significantly increased our understanding of human biology and disease. However, high costs and both logistic and technical challenges have limited its application to clinical studies^16^. In this context, a broader use of single-cell technologies could provide valuable insights to answer critical questions, such as the identification of molecular mechanisms associated with response and resistance^47–49^. Hence, ready-to-use commercial products that allow streamlined sample preservation and simplified logistics, such as the GEM-X Flex Gene Expression kit developed by 10X Genomics, hold the promise of lowering the bar for a broader application of scRNA-seq in clinical practice. In this regard, the possibility of using FFPE blocks can help address some of the limitations associated with the use of fresh or frozen material. First, fixing samples in formalin and embedding them in paraffin is a straightforward process, which is routinely used for pathology assessment, and it helps maintain the structural integrity and molecular composition of the tissue over time^23^. This sample preservation is particularly advantageous for retrospective studies, as it enables the analysis of samples collected years or even decades ago, thus unlocking valuable clinical and biological insights from precious archival collections and clinical cohorts. Moreover, the possibility of using consecutive sections from the FFPE blocks for standard histological analysis and scRNA-seq analysis allows direct comparison of the data, providing a more comprehensive understanding of the cellular composition, and gene expression profiles in the tissue. This dual approach enhances the accuracy of cell type identification and the mapping of cellular heterogeneity within the tissue and may help the clinical interpretation of the scRNA-seq data^50^.

In the current work we evaluate the performance of the GEM-X Flex protocol and its applicability to the clinical studies, benchmarking it against the standard Chromium Next GEM Single Cell 3’ and 5’ solutions in both frozen human tumors and FFPE tumor biopsies. Our analysis demonstrates that, in general, the GEM-X Flex protocol is more sensitive than the standard 10X Genomics protocols, allowing us to identify more genes and transcripts (UMIs) per cell. Ambient RNA contamination is a significant challenge when working with frozen samples analyzed by snRNA-seq, as it is usually the case of biopsies from multisite clinical trials^51^. This contamination can introduce noise into the data, making it difficult to distinguish true biological signals. Our data show that regardless of the pre-processing strategy, the tumor samples and FFPE blocks processed with the GEM-X Flex protocol retained more counts after ambient RNA removal using CellBender, suggesting a lower contamination with ambient RNA than the standard 10X Genomics solutions. These observations are in line with previous reports^12,45,52^ and can be attributed to a combination of general improvements in the GEM-X Flex protocol, such as the new probe-based chemistry and the advantages derived from tissue fixation. Of note, the probe-based approach also imposes some challenges and limitations as it cannot exclude that part of the signal is due to background noise (unspecific probe hybridization) and that the gene of interest could be missing, because not the whole transcriptome is sequenced. In summary, while the GEM-X Flex protocol presents some challenges, its enhanced sensitivity and reduced ambient RNA contribute to potential improvements in generating high-quality data and gaining valuable insights in clinical research.

The choice of easily implementable and standardized collection and preservation methods can help sample handling in the clinical setting^19^. However, different preservation strategies require the use of specific dissociation protocols that can lead to selective degradation of certain cell types. This phenomenon can result in biased cell population profiles that could influence the interpretation of the data, especially in heterogeneous tissues where rare cell types are crucial for understanding disease mechanisms^20,53^. In our analysis of frozen tumor samples, non-malignant cell subsets, including major cell types such as CD4, CD8, and NK cells, remain consistent across all protocols. However, significant differences are observed in the capture of less abundant and more fragile cell types, such as plasma cells, basophils, and fibroblasts, which are better preserved following the Chop/Fix protocol. This is also the case in the comparison of the clinical trial samples, in which the FFPE blocks processed with the GEM-X Flex protocol display a higher diversity, with more cell types represented. DCs, pDCs, and neutrophils are usually rare on standard scRNA-seq analysis and they are exclusively represented in the data derived from the FFPE blocks^54^. Nevertheless, we cannot fully exclude that our results are partially driven by the cellular representation in the original clinical material and not exclusively by the ability of the different workflows to preserve distinct cell types. Overall, these findings are consistent with previously published studies^41,45^ demonstrating how fixation protocols can help maintain the integrity of fragile cell types by preserving a more complex cellular composition.

We focused part of our analysis on T-cells, which are one of the main cell types of interest when analyzing biopsies of cancer patients, as they are the key effector of anti-tumor therapies such as CAR-T cells, check-point inhibitors or T-cell bispecifics^55,56^. However, our capacity to resolve and classify T-cells according to their activation status is strictly dependent on the quality of the samples, as it relies on the use of partially overlapping gene signatures^57,58^. When applied to FFPE samples, the GEM-X Flex protocol allowed us to assign most of the T-cells in the samples to the different functional subpopulations (naive, central memory, effector memory, cytotoxic, pre-dysfunctional, dysfunctional, regulatory, proliferating), which was not the case in the nuclei samples analyzed with the standard chemistry. This increase in resolution may be crucial in the context of clinical trials, as T-cell states and dynamics are often used to evaluate patient responses and therapeutic efficacy. However, TCR sequencing is not yet available in combination with the GEM-X Flex chemistry, so the use of the standard 10X Genomics 5’ chemistry or a similar solution from a different vendor is recommended for studies in which single cell TCR sequencing is required^59^.

We demonstrate that the use of FFPE samples with GEM-X Flex enables an enhanced detection of biologically relevant signals, linked to the mode of action of our drug candidate RO7119929. As previously described by Yoo et al.^44^, we further confirm the treatment-induced increase of inflammatory gene signatures, related to type I interferon response, viral sensing and antiviral IFN response signatures, which were identified as key pharmacodynamic markers, using bulk RNA-seq. Additionally, the analysis of matched baseline and on-treatment biopsies shows RO7119929-induced shift of immune cells such as a change in TAMs polarization, and the upregulation of interferon-stimulated genes and TLR7 downstream targets. These findings underscore the enhanced sensitivity of GEM-X Flex in detecting clinically relevant changes in gene expression signatures, highlighting its usability in biomarker discovery and translational research.

In summary, the improvements in data quality and sensitivity, together with the simplified logistics, streamlined lab protocols, and multiplexing capabilities offered by the GEM-X Flex technology, make it a suitable tool to overcome the key challenges for the application of scRNA-seq in clinical settings and may help increasing the use of this technology in clinical trials.

## Material & Methods

### Fresh lung carcinoma specimen

Fresh lung carcinoma specimen was sourced from Fidelis Research under the approved ethical guidelines. The fresh tumor sample was collected and transported (within 24 hr) in MACS tissue storage solution (Miltenyi, 130-100-008) at 4°C. Upon arrival, the specimen was manually dissected into small fragments (2×2 mm^2^), randomized, and up to 6-8 pieces were cryopreserved in pZerve freezing medium (Sigma, Z1653-60ML) at -80°C using a Corning CoolCell FTS30 (Corning, 43200). For long-term storage, cryovials were transferred to liquid nitrogen. The remaining tumor fragments were processed following the procedure summarized in **Figure 1A**.

### Clinical patient cohort and sample collection

As part of the Phase I clinical trial (NCT04338685) evaluating the therapeutic efficacy of RO7119929, paired tumor biopsies were collected from hepatic tumor lesions (n=3, primary tumor diagnosis: head and neck cancer (n=2) and colorectal cancer) using an 18-gauge core needle biopsy. Samples were obtained at baseline and on-treatment (Day 29) following administration of 5 mg RO7119929 in the first dose expansion cohort. One biopsy core was snap-frozen and used for single-nuclei RNA-seq, while the second core was used for histological assessment of tumor content and immune cell infiltration, performed by a board-certified pathologist.

### Cell composition analysis of lung carcinoma specimen by flow-cytometry

Cryopreserved lung carcinoma fragments were thawed using a water bath (37°C) and subsequently transferred into a petri dish (Greiner Bio-One, 633180), and washed once with cold PBS (Gibco, 14190-136). Subsequently, fragments were manually minced using a scalpel into smaller pieces and digested using an enzymatic mix [MACS Tissue Storage Solution (Miltenyi, 130-130-263); Accutase (Pan Biotech, P10-21100); Bovine serum albumin (Sigma-Aldrich, A9576); collagenase IV (Worthington, LS004188); DNase I Type 4 (Sigma-Aldrich, D5025); Hyaluronidase (Sigma-Aldrich, H6254)] for 30 min at 37°C on a thermo shaker (Thermo Fisher Scientific, SHKE4450). The resulting single-cell suspension was strained through a 70 µm cell strainer and washed (15 min, 300 xg at 4°C) twice with cold RPMI media (Pan Biotech; P04-17500), and subsequently with cold PBS. The cells were incubated in a V-bottom 96-well plate (Corning, 353263) with Zombie UV live/dead stain (Biolegend, 423108) for 10 min in the dark. Samples were washed with PBS and subsequently with FACS buffer [500 mL PBS; 3% FCS (Anprotec, AC-SM-0014Hi); 2 mM EDTA (Sigma-Aldrich, D2650)]. Human TruStain FcX Blocking Solutions (Biolegend, 422302) was added to the samples and incubated for 10 min in the dark. Subsequently, 25 µL of antibody mix (**Supplementary Table 1**) diluted in BD Horizon Brilliant Stain Buffer Plus (BD, 566385), was added to the cells and incubated for 15 min in the dark. Cells were washed twice with FACS buffer (5 min, 300 xg at 4°C) and resuspended in 100 µL FACS buffer for acquisition. The sample was measured on a Cytek Aurora (Cytek Biosciences) and analyzed using FlowJo software (v10.10.0, BD Life Sciences).

### Processing of fresh and snap-frozen lung carcinoma fragments into nuclei suspensions for snRNA-seq

To obtain single nuclei suspension, fresh and snap-frozen lung carcinoma fragments (3 pieces/condition) were dissociated using the Singulator 100 System (S2 Genomics) in NIC+ Nuclei Isolation cartridge (GC Biotech, 100-215-725). The pre-configured protocol for low-volume nuclei isolation was used. Protector RNAse Inhibitor (Roche, 3335402001) was added to the precooled NIC+ cartridge to prevent RNA degradation. The resulting nuclei suspension was transferred to a 15 mL conical centrifuge tube (Falcon, 352096) on ice and filtered through a 40 µm cell strainer (pluriSelect, 43-10040-40). The sample was subsequently centrifuged 300 g for 10 min at 4°C, and the pellet was resuspended in 500 µL of Nuclei Storage Buffer (GC Biotech, 100-215-725), supplemented with 0.2 U/µL RNase inhibitor. A final centrifugation step was performed under the same conditions. The nuclei suspension obtained from the fresh tissue was split into two parts. One part (**sample A**) was used for snRNA-seq analysis using the automated GEM-X Universal 5’ Gene Expression workflow on the Chromium Connect (10X Genomics, Protocol CG0002384 | Rev G). The second part of nuclei suspension from the fresh fragments (**sample B**) and the one derived from the snap-frozen fragments (**sample C**) were fixed according to the Fixation of Cells & Nuclei for Chromium Fixed RNA Profiling (10X Genomics, Protocol CG000478 | Rev C) for 24 hr at 4°C and then quenched. Fixed and quenched single-nuclei suspensions from Sample B and C were further processed for snRNA-seq analysis according to the Chromium Fixed RNA Profiling Reagent Kits User Guide (10X Genomics, Protocol CG000477 | Rev C). In all cases, the nuclei concentration of the samples was determined using a Nexcelom Cellometer Auto 2000 (Nexcelom Bioscience), and 10.000 nuclei were loaded into a 10X Genomics Chromium iX instrument. The concentration of the cDNAs and the final libraries was quantified using an Invitrogen Qubit 4 fluorometer (Thermo Fisher Scientific, Q33238), and the 1x dsDNA High Sensitivity Assay kit (Thermo Fisher Scientific, Q33231). The fragment size distribution was determined using a High Sensitivity DNA kit (Agilent, 5067-4626) on an Agilent 2100 Bioanalyzer automated electrophoresis instrument (Agilent). The libraries were sequenced in a NovaSeq6000 sequencer (Illumina Inc.) according to the corresponding 10X Genomics recommendations.

### Processing of fixed lung carcinoma fragments (Chop/Fix protocol)

To obtain single nuclei suspension from fixed lung carcinoma fragments (**sample D**), the samples were processed according to the Tissue Fixation & Dissociation Protocol for Chromium Fixed RNA Profiling (10X Genomics, Protocol CG000553 | Rev B). Briefly, tumor pieces were harvested and placed in a fixative solution, containing Fix & Perm Buffer (10X Genomics, PN 2000517) and 37% formaldehyde (Fisher BioReagents, BP531-25), to be manually dissected into small fragments as described above. Samples were fixed overnight at 4°C, washed twice in PBS, and finally kept in Quench Buffer (10X Genomics PN 2000516) on ice. Fixed and quenched samples were dissociated using Liberase TH (Millipore Sigma, 5401135001) mix and the gentleMACS Octo Dissociator with heaters (Miltenyi Biotec, 130-096-427). Dissociated samples were filtered through 30 µM Pre-Separation Filters (Miltenyi, 130-041-407), washed with PBS, and resuspended in chilled Quench Buffer. The concentration was determined using a Nexcelom Cellometer Auto 2000, and 10,000 nuclei were loaded into a 10X Genomics Chromium iX instrument. Cell capture and library preparation were performed according to the Chromium Fixed RNA Profiling Reagent Kits User Guide (10X Genomics, Protocol CG000477 | Rev C). The concentration of the final library was quantified using the Invitrogen Qubit 4 fluorometer, and the 1x dsDNA High Sensitivity Assay kit. The fragment size distribution was determined using an Agilent 2100 Bioanalyzer automated electrophoresis instrument (High Sensitivity DNA kit). The libraries were finally sequenced in a NovaSeq6000 sequencer (Illumina Inc.) according to 10X Genomics recommendations.

### Single-nuclei RNA-seq from tissue samples collected during the clinical study

For the clinical trial analysis, single-nuclei RNA-seq was performed on snap-frozen biopsies using the 10X Genomics Chromium Next GEM Automated Single Cell 3ʹ Reagent Kits v3.1, following the manufacturer’s protocol. Briefly, 10,000 nuclei per sample, diluted at a density of 300–2,500 cells/μL in PBS plus 0.04% BSA (determined by Cellometer Auto 2000 (Nexelom Bioscience)), were targeted. All samples were run in triplicates. The quality and concentration of both cDNA and libraries were assessed using an Agilent BioAnalyzer with High Sensitivity kit (both Agilent) and Qubit Fluorometer with 1x dsDNA High Sensitivity Assay kit according to the manufacturer’s recommendation. For sequencing, samples were mixed equimolecularly and sequenced on a HiSeq4000 (Illumina Inc.) following 10X Genomics recommendations.

### Processing of clinical FFPE tissue samples for scRNA-seq

For the present study, we accessed the leftover FFPE tissue blocks from the baseline and on-treatment biopsies of three patients from the NCT04338685 trial. The FFPE blocks were processed according to the Sample Preparation from FFPE Tissue Sections for Chromium Fixed RNA Profiling (10X Genomics, Protocol CG000632 | Rev D), to generate a single-cell suspension. Briefly, tissue biopsies were cut out of the FFPE block, and paraffińs excess was manually removed with a scalpel. The tissue samples were deparaffinized by washing them at room temperature (RT) in xylene (Sigma Aldrich, 214736-1L) three times for 10 min, and progressively rehydrated by incubating them for 30 s at RT in decreasing ethanol (Roth, 5054.1) concentration (twice in 100% ethanol, once in 70% ethanol, and once in 50% ethanol). Finally, the samples were washed for 30 s at RT in distilled water and kept on ice in PBS. The tissue biopsies were dissociated according to the gentleMACS Octo Dissociator for FFPE Tissue protocol, using Liberase TH (Millipore Sigma, 5401135001) mix and the gentleMACS Octo Dissociator with heaters. The obtained single-cell suspension was filtered through 30 µM Pre-Separation Filters, placed on ice, washed twice with ice-cold PBS, and resuspended in Quench Buffer. Cells were counted with Cellometer Auto 2000 (Nexcelom Bioscience). Single nuclei suspensions were further processed for snRNA-seq analysis according to the Chromium Fixed RNA Profiling Reagent Kits User Guide (10X Genomics, Protocol CG000477 | Rev C). The concentration of the final library was quantified using an Invitrogen Qubit 4 fluorometer, and the 1x dsDNA High Sensitivity Assay kit. The fragment size distribution was determined using an Agilent 2100 Bioanalyzer automated electrophoresis instrument (High Sensitivity DNA kit). The libraries were sequenced in an Illumina NovaSeq6000 sequencer according to 10X Genomics recommendations.

### Bioinformatic analysis

Sequencing files were converted to FASTQ files using the Cell Ranger pipeline and subsequently aligned using CellRanger count v7.1.0 to either the human transcriptome (GRCh38), for the standard 5’ chemistry samples, or to the human Probe Set (v1.0.1), in the case of Flex samples. Firstly, an independent analysis was performed per chemistry (5’ or Flex) and dataset (namely “experimental conditions” or “clinical study”) to leverage each chemistry’s characteristics for filtering and annotation purposes (e.g so as not to restrict the nuclei samples to probe set genes). For this, the raw count matrices were aggregated into 4 data objects and processed with the besca standard workflow (besca v2.5 and scanpy v1.9.1)^60^. A first filtering was performed with the parameters min_counts=1000; min_genes=500; min_cells=30; n_genes=12000 for Flex; 8000 for 5’; max_counts=200000 for Flex; 50000 for single-cell 5’; 150000 for nuclei 5’; percent_mito=5% for Flex; 1.5% for nuclei 5’. Ambient RNA was then removed by running CellBender v0.2.2 with per sample expected-cells, total-droplets and low-count-threshold parameters based on the Cell Ranger UMI barcode rank plots. The uncorrected count matrices were replaced with the corrected ones in the data object and cells with <100 genes, <200 counts or called empty by CellBender were further removed. One sample from the clinical study dataset (nuclei_patient3_Treated) failed QC and was removed for downstream analysis, mainly due to very low percentage of reads mapped confidently to transcriptome, which resulted in very low per-cell UMI counts and almost 80% filtered cells. After filtering and ambient correction, counts were normalized to log-transformed UMI counts per 10,000 reads [log(CP10K+1)], and the most variable genes (having a minimum mean expression of 0.0125, a maximum mean expression of 3 and a minimum dispersion of 0.5) were used for principal component analysis (PCA). The first 50 PCs were used for calculating the 10 nearest neighbors and the resulting neighborhood graph was used for Leiden clustering (resolution=2) as well as for UMAP-based visualization.

Cell type annotation was initially performed using the Sig-annot semi-automated Besca module, a signature-based hierarchical cell annotation method^60^, followed by intense manual inspection. The used human signatures, configuration and nomenclature files are in https://github.com/bedapub/besca/tree/master/besca/datasets. The cluster-based annotations were further refined at cell level using a combination of per-cell scanpy signature scores to e.g. resolve mixed clusters or annotate doublets. Cells were annotated at different levels of granularity (0: broad annotation; 1: main cell types; 2: including cell subtypes) following Besca hierarchy. To further identify doublets, scDblFinder v1.14.0 was also run per sample with default parameters.

To generate figures for this manuscript, each dataset was re-analyzed with all samples together (both Flex and 5’ chemistries) using the ambient corrected counts, and the previously filtered and annotated cells. If not otherwise stated, the genes were restricted to the probeset. UMAP-based visualization was run either on unintegrated data or on integrated data by using the BBKNN method, using chemistry as the batch variable.

For the tissue dissociation stress analysis, we used the following gene signature: *SRSF5, FOS, GADD45G, HES1, PIM1, ATF3, CCN1, ACTB, TENT5A, GJA1, JUN, ACTG1* adapted from Neuschulz 2023^46^ (Supplementary Table EV3). The per-cell gene signature expression score was calculated separately for each sample using the Scanpy score_genes() function with default parameters. Then, the score was normalized to min=0, max=1 (formula [x - x.min()] / [x.max()-x.min()], where x is the cell score), and distributions were visualized and compared in violin with embedded boxplots.

For the dedicated T-cell states analysis, T-cells were reanalyzed and annotated separately (nuclei 3’ and FFPE samples) by manually attributing clusters to naive, central memory, effector memory, cytotoxic, pre-dysfunctional, dysfunctional, regulatory, proliferating or unresolved T-cells using a combination of Besca and Leun signatures^57^, as shown in **Supplementary Figure 3C**. Samples with less than 30 cells were excluded from this analysis (only 2 nuclei samples left).

For the macrophage polarization analysis, macrophages were selected and the per-cell Scanpy score was calculated per chemistry with the score_genes() function and default parameters for the following myeloid polarization signatures: M1-like (CXCL10, CXCL11, CD274, CD300E, IDO1, ISG15) and M2-like (MRC2, CXCL12, CD163, MRC’, MSR1, TREM2, CD163L1). Subsequently, a per sample average score was calculated and used for plotting.

## Ethics approval

The study was approved by all applicable independent Ethics Committees and Regulatory Authorities and was conducted in compliance with the Declaration of Helsinki, ICH/Good Clinical Practice E6(R1), European Union Clinical Trials Directive, and local regulatory requirements. The study was registered on ClinicalTrials.gov (NCT04338685). Written informed consent was obtained from all participants before the assessment of eligibility.

## Availability of data and materials

The individual patient-level data generated and/or analysed during the current study are not publicly available due to Roche company policy and patient privacy reasons. Qualified researchers may request access to individual patient-level data through the clinical study data request platform (https://vivli.org/). Further details on Roche’s criteria for eligible studies are available here (https://vivli.org/ourmember/roche/). For further details on Roche’s Global Policy on the Sharing of Clinical Information and how to request access to related clinical study documents, see here (https://www.roche.com/innovation/process/clinical-trials/data-sharing/).

## Authors’ contributions

M.A., E.Y., and S.D conceptualised the study. M.A. and K.P. conducted the research investigation process, specifically performing the experiments, and data collection.. L.A-S. performed formal bioinformatics analyses. M.A., L.A-S., S.S., E.Y., and S.D. interpreted experimental and sequencing data. S.S. and S.N. advised regarding bioinformatics analyses. C.Y. generated flow-cytometry data. T.H. advised regarding wet-lab experiments. M.A.C. provided clinical trial resources and supported data interpretation. M.B. and S.D. supervised the research activity. M.A., L.A-S., E.Y., and S.D. wrote the manuscript. All authors reviewed and approved of the manuscript.

## Acknowledgment

We sincerely acknowledge Vera Griesser, Desiree Von Tell, and Kim Schneider from the Genomics 360 Lab at Roche for their support with sequencing. We also extend our gratitude to Natascha Rieder for her expert pathological evaluation of the tissue sections. Additionally, we deeply appreciate the invaluable support of Naailah Roheemun, Quincy Dekempe, Monika Friedrich, Romi Federson, and Gordon Heidkamp in establishing the 10X Genomics GEM-X Flex protocol, while providing access to essential equipment, which greatly facilitated this work. Finally, we thank the entire pRED project team, as well as the clinical and biomarker teams involved in the NCT04338685 trial, and Sabine Hoves for proofreading the manuscript.

