## Supplemental Figures and Table for "Performance of the 10X Genomics Flex Single-Cell Sequencing Assay and its Application to Overcome Challenges in Clinical Trial Samples"

**Short title: Overcoming Clinical Trial Challenges with 10X Genomics Flex Sequencing**


##### Authors

**Martina Antoniolli[1], *Llucia Albertí Servera[2], Kerstin Paetzold[1], Stephan Schmeing[2], Carmen Yong[3], Sina Nassiri[2], Tamara Hüsser[3], Michael A. Cannarile[1], Marina Bacac[3], #Emilio Yángüez[3], #Steffen Dettling[2]*

##### Affiliations
**1** Roche Innovation Center Munich, Roche Pharmaceutical Research and Early Development (pRED), Penzberg, Germany
**2** Roche Innovation Center Basel, pRED, Basel, Switzerland
**3** Roche Innovation Center Zurich, pRED, Schlieren, Switzerland

* shared first authorship
### shared last authorship

**Corresponding author**: Dr. Steffen Dettling, Roche Innovation Center Basel, Roche Pharmaceutical Research and Early Development (pRED), Grenzacherstrasse 124, 4058 Basel, Switzerland. Tel. +41 79 893 76 34, E-Mail

### Additional Files

**Additional file 1.doc - All supplementary files for this study**

- **Supplementary Figure 1.** Extended data from the quality control metrics of the tumor samples.
- **Supplementary Figure 2.** Extended data from the cell annotation and quantification of tumor samples.
- **Supplementary Figure 3.** Extended data from the analysis of the clinical samples.
- **Supplementary Table 1.** Antibodies used for flow-cytometry

**A** **B** **C**

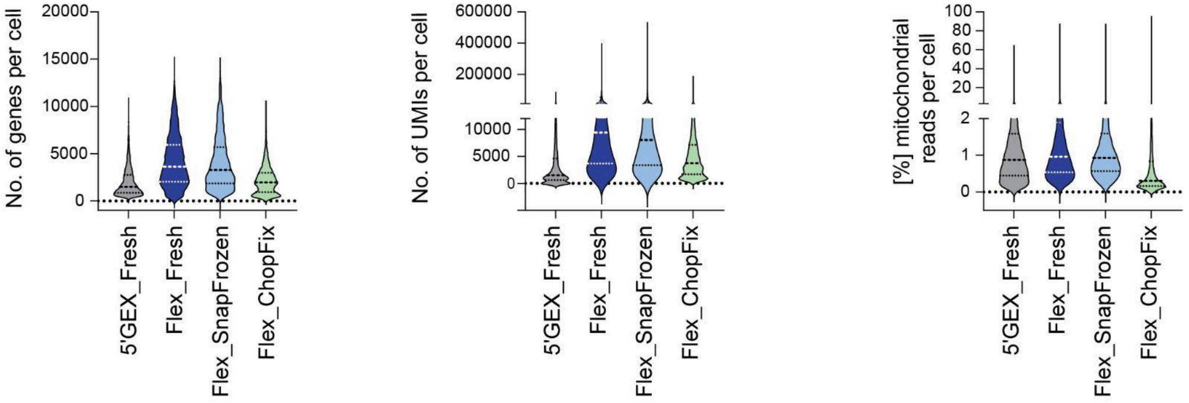


**D**

| | [%] Mito | | No. genes | | No. UMI | |
|---|---|---|---|---|---|---|
| | Probeset | ALL | Probeset | ALL | Probeset | ALL |
| **5´GEX_Fresh** | 0.74 | 0.88 | 1337 | 1497 | 1954 | 2258 |
| **Flex_Fresh** | 0.96 | | 3634 | | 9401 | |
| **Flex_SnapFrozen** | 0.93 | | 3274 | | 8019 | |
| **Flex_ChopFix** | 0.30 | | 1966 | | 3730 | |

**F**

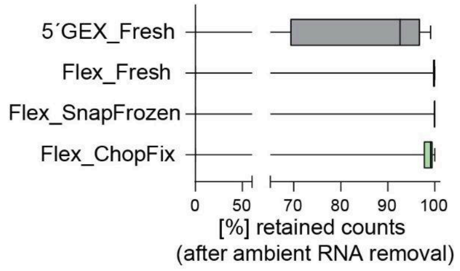


**E**

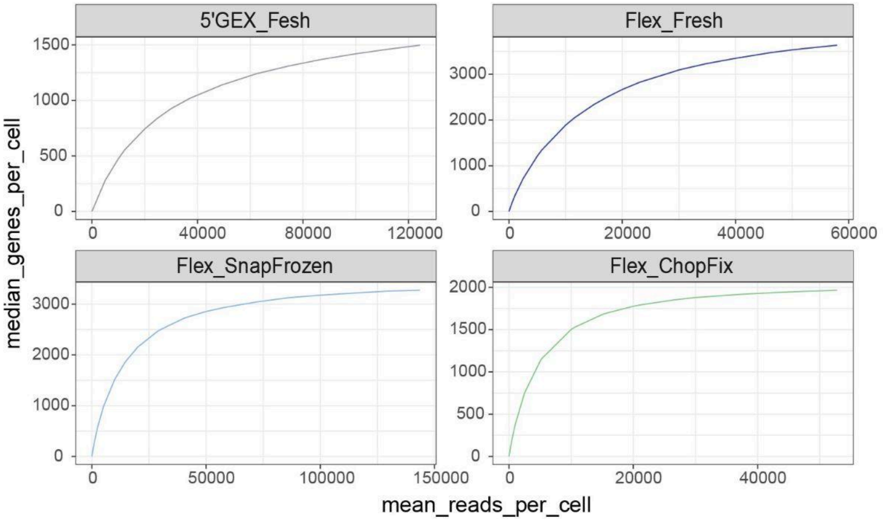


**Supplementary Figure 1. Extended data from the quality control metrics of the tumor samples. (A-C)** Distribution of the number of genes **(A)**, the number of Unique Molecular

Identifiers (UMIs, **B**), or the percentage of reads mapping to mitochondrial genes **(C)** per nuclei/cells detected in the samples processed with the different protocols (non-filtered CellRanger barcodes, all detected genes). **(D)** Summary table with the median values for the QC metrics in A-C and per sample. For the 5' GEX sample, the median is computed with either the genes present in the GEM-X Flex probeset only, or with all genes. **(E)** Number of median genes detected per nuclei/cells at different sequencing depths for all the samples. **(F)** Distribution of the number of counts per nuclei/cells retained after ambient RNA removal using CellBender in the different samples (all CellRanger nuclei/cells are represented).

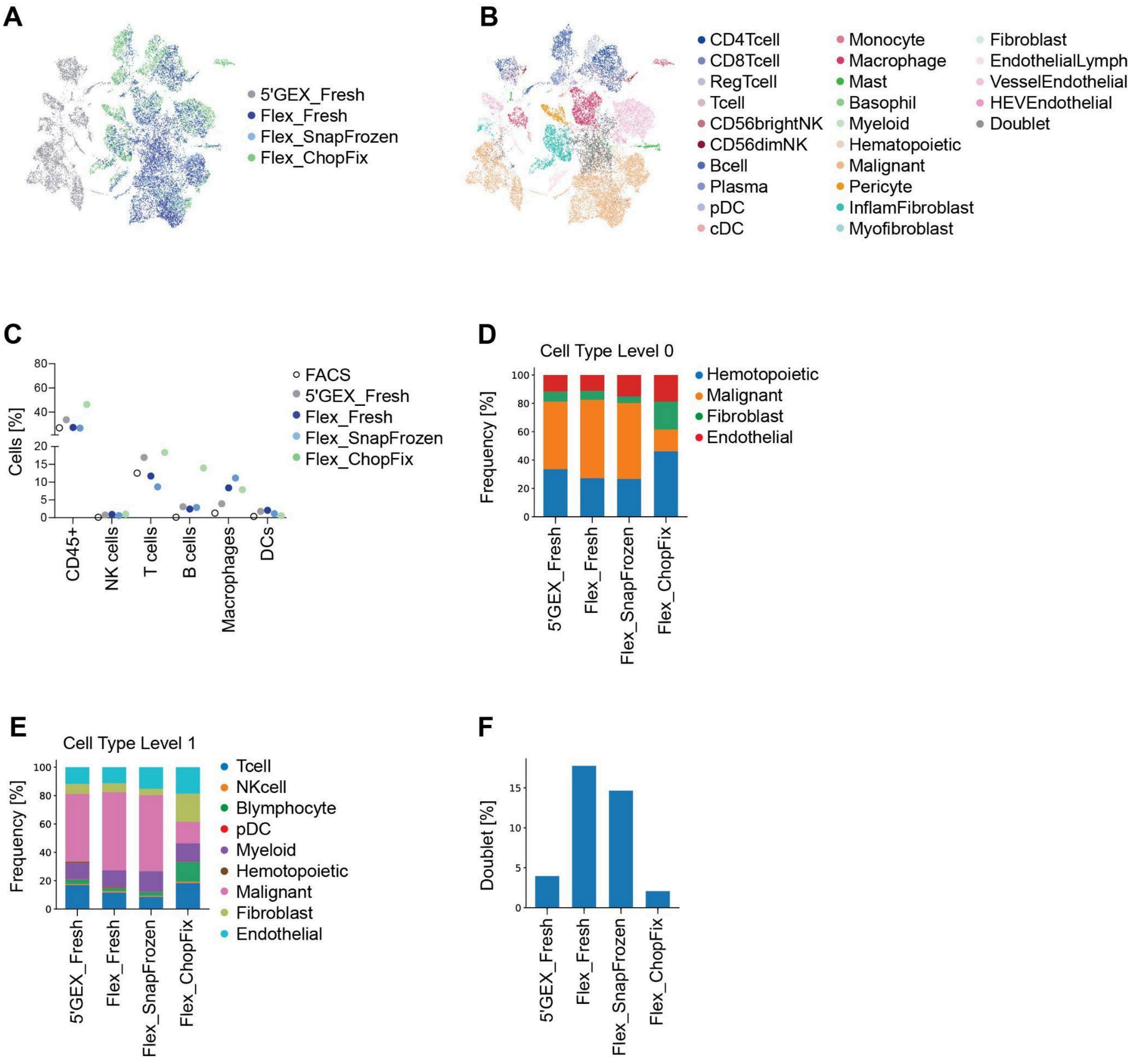


**Supplementary Figure 2. Extended data from the cell annotation and quantification of tumor samples. (A)** Uniform manifold approximation and projection (UMAP) plot of unintegrated data, colored by experimental condition. **(B)** UMAP plot of unintegrated data, colored by cell type (granular annotation level 2). **(C)** Cell type abundance of selected cell types represented as a percentage of the total number of nuclei/cells identified by sequencing (using the different protocols) or by fluorescence-activated cell sorter (FACS) from the matching sample as a reference. **(D-E)** Cell type distribution as percentage of the total number of nuclei/cells identified in the different samples with broad annotation (level 0, **D**) or intermediate annotation granularity (level 1, **E**). **(F)** Percentage of final doublets in the samples processed with the different chemistries.

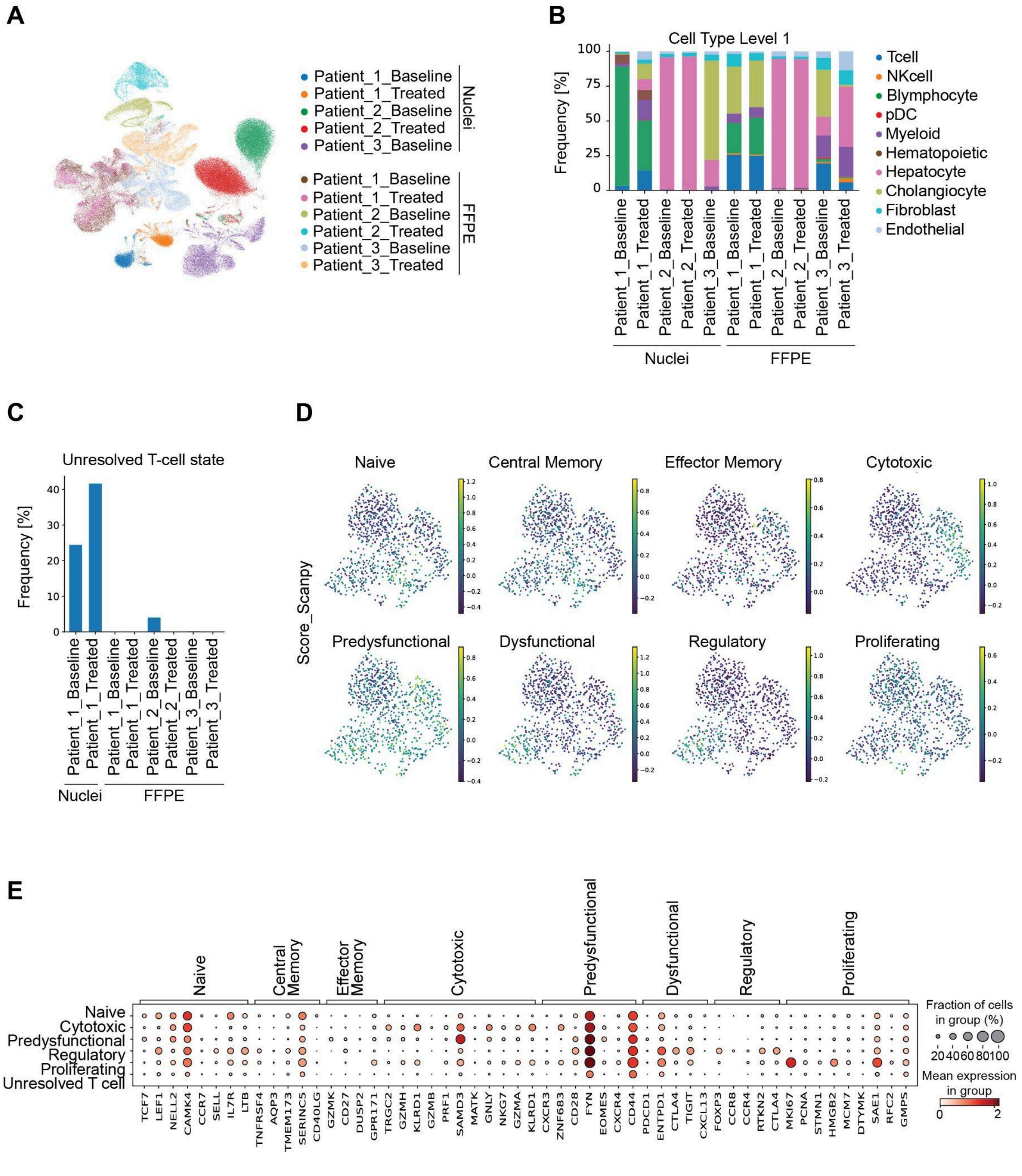


**Supplementary Figure 3. Extended data from the analysis of the clinical samples. (A)** Uniform manifold approximation and projection (UMAP) plot of unintegrated data, colored by sample. **(B)** Cell type distribution as percentage of the total nuclei/cells identified in the different samples (annotation level 1). **(C)** Percentage of T-cells per sample for which the T-cell state could not be resolved. **(D)** UMAP plot with T-cells from nuclei samples colored according to the gene signature scores for the indicated T-cell states. **(E)** Dotplot with T-cells (stratified per annotated state) from nuclei samples showing the gene expression level (color) and fraction of positive cells (dot size) for the marker genes of each T-cell state signature.

#### Supplementary Table 1. Antibodies used for flow-cytometry

| Marker | Fluorochrome | Supplier | Cat # | Clone | Final dilution |
|---|---|---|---|---|---|
| CCR7 | BUV661 | BD | 749824 | 2-L1-A | 100 |
| CD11B | BB515 | BD | 564454 | M1/70 | 100 |
| CD11c | PerCP-eFluor 710 | eBioscience | 46-0116-42 | 3.9 | 100 |
| CD123 | APC/Fire 810 | Biolegend | 306053 | 6H6 | 100 |
| CD14 | PerCP | Biolegend | 325631 | HCD14 | 200 |
| CD163 | PE-Dazzle 594 | Biolegend | 333623 | GH1/61 | 100 |
| CD19 | Alexa Fluor 532 | eBioscience | 58-0199-42 | HIB19 | 200 |
| CD218a | APC | Biolegend | 313813 | H44 | 100 |
| CD25 | BUV395 | BD | 564034 | 2A3 | 100 |
| CD28 | BUV563 | BD | 748476 | L293 | 100 |
| CD3 | BV570 | Biolegend | 300436 | UCHT1 | 200 |
| CD304 | BV510 | Biolegend | 354515 | 12C2 | 100 |
| CD31 | BV605 | Biolegend | 303122 | WM59 | 100 |
| CD39 | BV785 | Biolegend | 328239 | A1 | 100 |
| CD4 | SparkYG593 | Biolegend | 344672 | SK3 | 100 |
| CD45 | BUV805 | BD | 612891 | HI30 | 150 |
| CD45RA | BV750 | Biolegend | 304165 | HI100 | 150 |
| CD56 | PE-Fire 640 | Biolegend | 392431 | QA17A16 | 200 |
| CD8a | BUV615 | BD | 751518 | RPA-T8 | 200 |
| CD90 | BV650 | Biolegend | 328144 | 5 E10 | 100 |
| Fixable Viability Dye | Zombie UV | Biolegend | 423108 | | 2000 |
| HLA-DR/DP/DQ | BUV496 | BD | 741157 | Tu39 | 100 |
| LAG-3 | PE-Cy5 | Biolegend | 369345 | 11C3C65 | 100 |
| PD-1 | PE-Fire700 | Biolegend | 621621 | A17188B | 100 |
| PD-L1 | PE | Biolegend | 329706 | 29E.2A3 | 100 |
| Tim-3 | PE-Fire810 | Biolegend | 345059 | F38-2E2 | 100 |
